# Repeatability and reproducibility of the *wzi* High Resolution Melting-based clustering analysis for *Klebsiella pneumoniae* typing

**DOI:** 10.1101/2020.06.19.161703

**Authors:** Ajay Ratan Pasala, Matteo Perini, Aurora Piazza, Simona Panelli, Domenico Di Carlo, Cristian Loretelli, Alessandra Cafiso, Sonia Inglese, Gian Vincenzo Zuccotti, Francesco Comandatore

## Abstract

**Aims:** High Resolution Melting (HRM) is a fast closed-tube method for nucleotide variant scanning applicable for bacterial species identification or molecular typing. Recently a novel HRM-based method for *Klebsiella pneumoniae* typing has been proposed: it consists of an HRM protocol designed on the capsular *wzi* gene and an HRM-based algorithm of strains clustering. In this study, we evaluated the repeatability and reproducibility of this method.

**Methods and Results:** We performed HRM typing of a set of *K. pneumoniae* strains, on three different instruments and by two different operators. The results showed that operators do not affect measured melting temperatures while different instruments can. Despite this, we found that strains clustering analysis remains perfectly consistent performing the analysis using the MeltingPlot tool separately on the data from the three instruments.

**Conclusions:** The HRM protocol under study was highly repeatable and thus reliable for large studies, even when several operators are involved.

**Significance and Impact of the Study:** Our results show that the *wzi*-HRM protocol for *K. pneumoniae* typing is suitable for multicenter studies, even if different instruments are used.

## Introduction

*Klebsiella pneumoniae* is a Gram-negative opportunistic pathogen often present in the gut of healthy individuals but also able to cause severe infections. Furthermore, the bacterium is one of the most important nosocomial pathogens, causing healthcare-acquired infections worldwide with a mortality rate ranging from 20 to 70% (Angus et al., 2001; Mayr, Yende and Angus, 2014). Indeed, *K. pneumoniae* has been described as an “urgent threat to human health” by the United States Centers for Disease Control and Prevention (CDC) and the World Health Organization (WHO) (Munoz-Price et al., 2013). Genomic studies revealed that, despite the high genetic variability of the bacterium (Gaiarsa et al., 2015; Holt et al., 2015), most of the nosocomial outbreaks are caused by only few Multi Drug Resistant (MDR) clones, in particular ST258, ST512, ST307, ST11, ST101 and ST15 (David et al., 2019; Wyres et al., 2019). Thus, a genetic-based nosocomial surveillance can represent an important tool to promptly detect *K. pneumoniae* high risk clones in the hospital setting.

High Resolution Melting (HRM)-based typing is a promising tool for clinical and epidemiological applications (Tamburro and Ripabelli, 2017). HRM is a fast closed-tube method to discriminate nucleotide variants on the basis of PCR amplicon melting temperature. The method is particularly reliable for nosocomial surveillance: it can be performed on most real-time PCR instruments; the entire protocol takes ∼5 hours and it is inexpensive (∼5$ per sample).

Perini and colleagues (Perini, Piazza, et al., 2020) proposed an HRM-based method for *K. pneumoniae* typing. The method consists of an HRM protocol designed on the hypervariable capsular gene *wzi* and followed by a strains clustering analysis based on the melting temperatures. The method was able to discriminate most of the *K. pneumoniae* Sequence Types (STs) known as “high risk” (Perini, Piazza, et al., 2020).

Different real-time PCR/HRM instruments can vary in thermal precision and melting temperature acquisition rate (Wittwer, 2009; Li et al., 2014). In literature, studies on HRM protocols designed for human samples revealed that the measured melting temperature can vary among the instruments (Wittwer, 2009; Li et al., 2014). In this study, we evaluated the repeatability and reproducibility (Schulten et al., 2000; Bustin et al., 2009) of the HRM method described by Perini and colleagues (Perini, Piazza, et al., 2020) repeating HRM typing on three different instruments by two operators.

## Material and Methods

### Dataset selection

The dataset for the analyses was a subset of the 82-strains collection analyzed by Perini and colleagues (Perini, Piazza, et al., 2020). In order to obtain an unbiased collection, we selected the strains considering the HRM clusters found by Perini and colleagues (Perini, Piazza, et al., 2020) and the *wzi* allele of the strains. More in detail, the strains were selected as follows:

- one strain was retrieved from each of the three HRM clusters containing one or two strains
- two strains were retrieved from each of the three clusters containing more than two strains
- three more strains were selected from the Cluster “ST11/ST101-like” (Perini, Piazza, *et* al., 2020) which contain many *wzi* alleles associated to epidemiologically relevant clades

A total of 14 *K. pneumoniae* strains belonging to eight different STs, including the highly epidemiologically relevant ST258, ST512, ST307, ST11, ST101 and ST15 (David et al., 2019) were selected (see Table S1 for details).

### Real-time PCR/HRM instruments

HRM analysis (see below) was performed on three different real-time PCR/HRM instruments:

- Bio-Rad CFX96 real-time PCR machine (Bio-Rad Laboratories), from here “CFX96”
- Eco Real-Time PCR System (Illumina), from here “Eco_RT”
- QuantStudio 6 Flex Real-Time PCR System (Applied Biosystems), from here “QS_6Flex”

The three instruments were placed in three different laboratories in two cities, for details see Table S2.

### DNA extraction

Bacterial strains were freshly streaked on MacConkey agar plate and incubated overnight at 37°C; then a single colony was inoculated into 5 mL of LB broth (DifcoTM) and incubated overnight at 37°C with vigorous shaking. For each strain, 1 × 109 cells have been used as starting material for total DNA extraction using the DNeasy blood and tissue kit (Qiagen) following the manufacturer’s instructions.

### High Resolution Melting analysis

For each strain, the extracted DNA was subjected to six HRM analyses: two operators (MP and AP) independently performed the HRM analyses on the three real-time PCR/HRM instruments listed above. In each of the six HRM analysis, three technical replicates were performed for each strain, amplified with the two primer pairs in the Perini and colleagues (Perini, Piazza, et al., 2020) HRM protocol (*wzi-3* and *wzi-4*). Negative controls were added in every HRM analysis for each primer pair.

The HRM reaction mix (10µl) contained: 5µl of 2x SsoAdvanced Universal SYBR® Green Supermix (BioRad, Hercules, California), 0.4µl of each primer (0.4µM) and 1µl of template DNA (25-50ng/µl). The thermal profile was as follows: 98°C for 2min, 40 cycles of [95°C for 7s, 61°C for 7s, and 72°C for 15s], 95°C for 2min, followed by HRM ramping from 70–95°C. Fluorescence data were acquired at increments of 0.5°C for CFX96, 0.3°C QuantStudio 6 Flex, and 0.1°C for Eco Real-Time PCR System. Each CFX96 and QuantStudio 6 Flex HRM analysis was performed in a single 96-well optical plate, while for the Eco Real-Time PCR System each HRM analysis required three 48-well optical plates.

DNA and reagents aliquots for all the experiments were prepared in advance to reduce the risk of contamination. In each experiment, the two operators independently prepared the HRM mixes in a pre-PCR ‘clean’ room using the same pipettes each day for each individual experiment.

### Statistical analysis

For each strain, the average of the melting temperatures (aTm) obtained from the three technical replicates were computed for *wzi-3* and *wzi-4* primer sets. A preliminary qualitative comparison of the aTms obtained by the different instruments and operators was performed reporting the median, minimum and maximum temperature differences for *wzi-3* and *wzi-4* aTms and plotting the aTm distributions by boxplots.

Then, the effects of operators or instruments on *wzi-3* and *wzi-4* aTms were investigated as independent and as combined factors. The statistical analyses were performed on 1,000,000 bootstrapping strain subsets, randomly selected with replacement. For each subset, aTms were analyzed using R v.3.6.1 (https://www.r-project.org/) as follows:

1. Independent factors:
  - Normality distribution of aTm values was tested by the Shapiro-Wilk test.
  - Homoskedasticity variances of aTm values between operators and among the three instruments were compared using F test and Bartlett test, respectively.
  - If aTms were normally distributed, the operators were compared using t-test (applying the Welch approximation in case of heteroskedasticity variance), otherwise using Mann-Whitney test.
  - If aTms were normally distributed, the instruments were compared using one-way ANOVA (ANalysis Of VAriance, or Welch one-way ANOVA in case of heteroskedasticity variance), otherwise Kruskal-Wallis test.
2. Combined factors:
  - The effects of operator and instrument, and their interactions, were evaluated using the non-parametric analysis of variance implemented in the art function of ARTool R 3.6.1 package (Kay, no date).

Then, we evaluated the percentage of subsets for which the effect of operator, instrument or their interaction were significant (p-value < 0.05).

### HRM clustering analysis

HRM data obtained for each instrument (CFX96, Eco_RT and QS_Flex6) were independently subjected to HRM clustering analysis using the MeltingPlot tool (Perini, Batisti Biffignandi, *et* al., 2020) (available at https://skynet.unimi.it/index.php/tools/meltingplot/). More in detail, for each strain the tool computes the replicates average melting temperatures (aTms) for each primer set. Using the igraph R library (http://igraph.org/), the tool builds a graph connecting the strains with aTm distance <= 0.5 °C for every primer set, and it clusters the strains on the basis of their betweenness (Perini, Batisti Biffignandi, et al., 2020).

## Results

### High Resolution Melting analysis

Fourteen *Klebsiella pneumoniae* strains were subjected to the HRM protocol proposed by Perini and colleagues (Perini, Piazza, et al., 2020) using *wzi-3* and *wzi-4* primer sets. The analysis was repeated by two operators on three different instruments, for a total of six experiments. The resulting average melting temperatures (aTms) are reported in Table S3.

### Statistical analysis

Boxplots of the aTms from the six HRM experiments are reported in Figure 1. The median, minimum and maximum aTm differences among operators/instruments are reported in Supplementary Table 4, for a total of 15 operator/instrument combinations per primer set.

**Figure 1:**
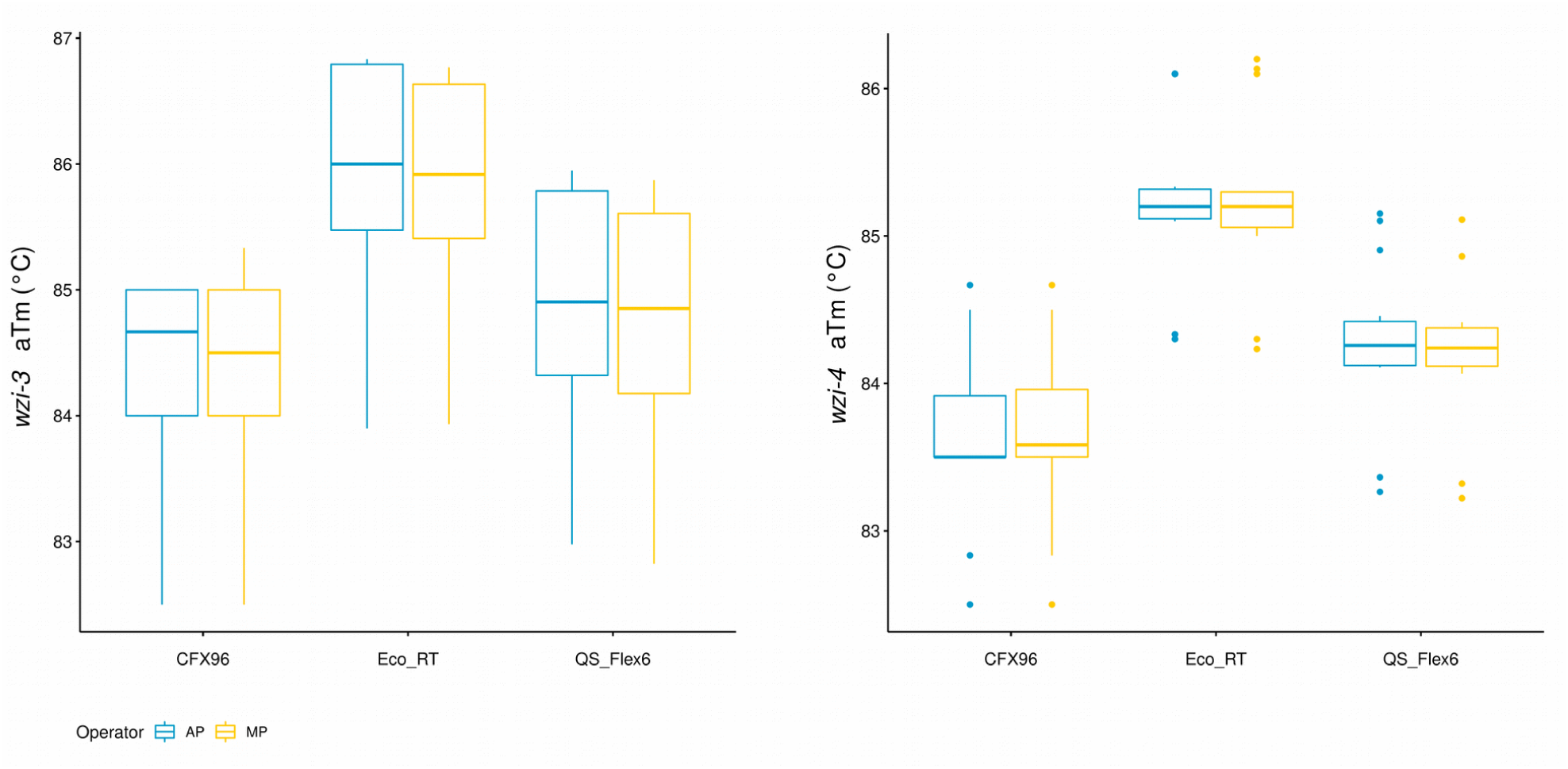
Boxplot of the *wzi-3* and *wzi-4* average melting temperatures. The distributions of wzi-3 and wzi-4 average melting temperatures (aTm) obtained by each instrument (“CFX96” for Biorad CFX96; “Eco_RT” for Illumina Eco Real-Time; “QS_Flex6” for Applied Biosystem QuantStudio 6 Flex) and operator (AP and MP; colored in blue and yellow, respectively) are shown. Boxes range between the 25th and the 75th quartiles, and bold horizontal lines represent the median values.

For either primer couples, the combinations of different operators on the same machine gave a maximum difference below 0.5 °C, the threshold set by Perini and colleagues (Perini, Piazza, *et* al., 2020) for clustering analysis. Conversely, all the combinations among different machines gave maximum differences above 0.5 °C.

The results of the statistical analysis are summarized in Table S5. For both *wzi-3* and *wzi-4*, T-test/Wilcox tests on the operators resulted non-significant (p-value >= 0.05) for the 100% of the 1,000,000 bootstrap replicates, while the instrument resulted significant (p-value < 0.05) for all of them. For *wzi-3*, the non-parametric analysis of the variance found the operator to be significant for 3.16% of the bootstraps replicates, the machine for 100% of replicates, and the interaction among the two factors (operator/instrument) was found to be significant for 0.64% of the bootstrap replicates. For *wzi-4*, the operator was found significant for 0.02% of the bootstraps replicates, the machine for 100% and the interaction for 0.78%.

### HRM clustering analysis

For each of the three instruments (CFX96, Eco_RT and QS_6Flex), the 14 strains were clustered on the basis of *wzi-3* and *wzi-4* aTms using the MeltingPlot tool (Perini, Batisti Biffignandi, *et* al., 2020). The results are reported in Figure 2 and Table 1. For each instrument, the clustering analysis grouped the 14 strains in six clusters and no strains were classified as “undetermined”. The strains clusters resulted totally conserved among the instruments.(Perini, Batisti Biffignandi, et al., 2020)

**Table 1.**
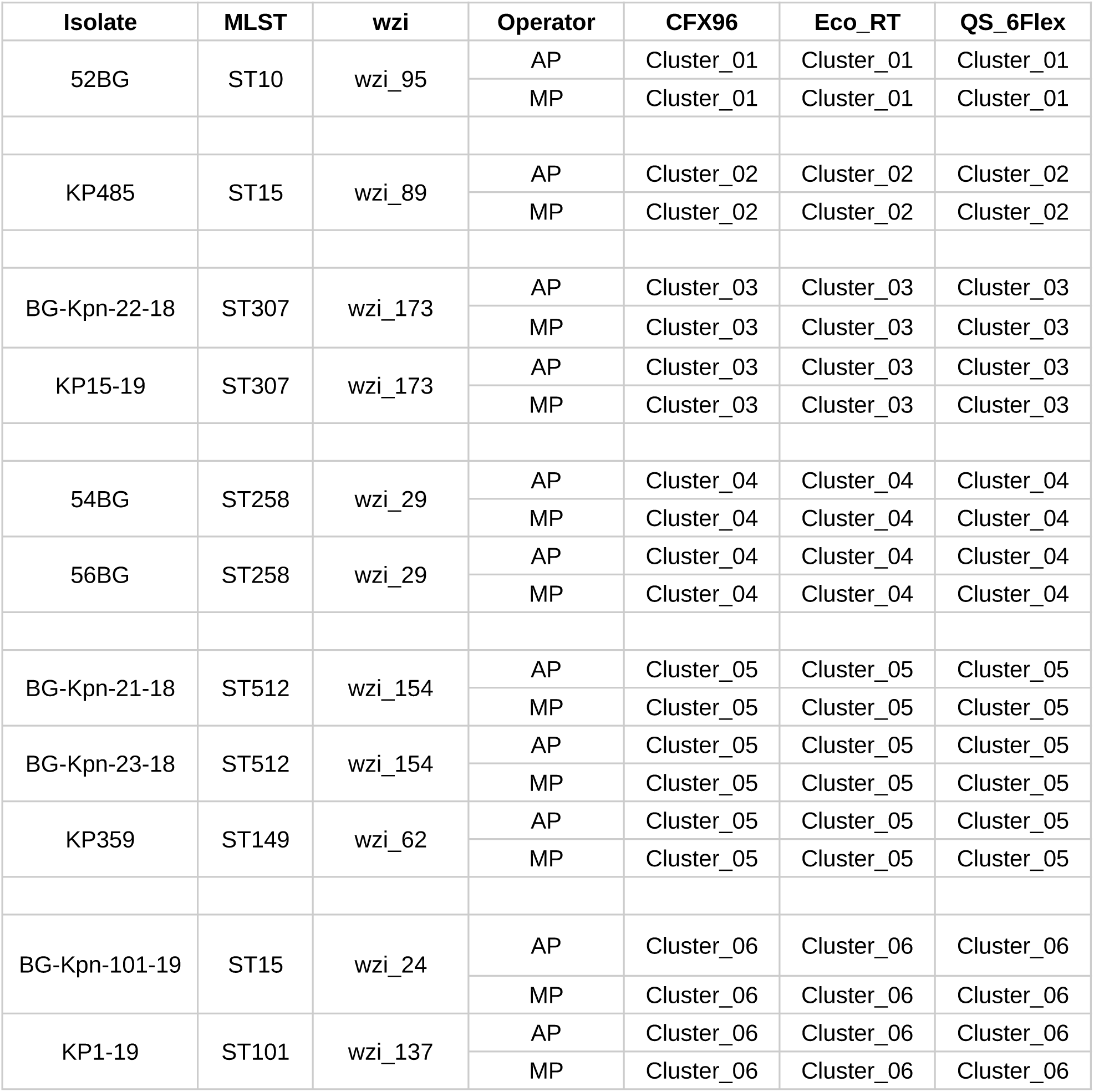
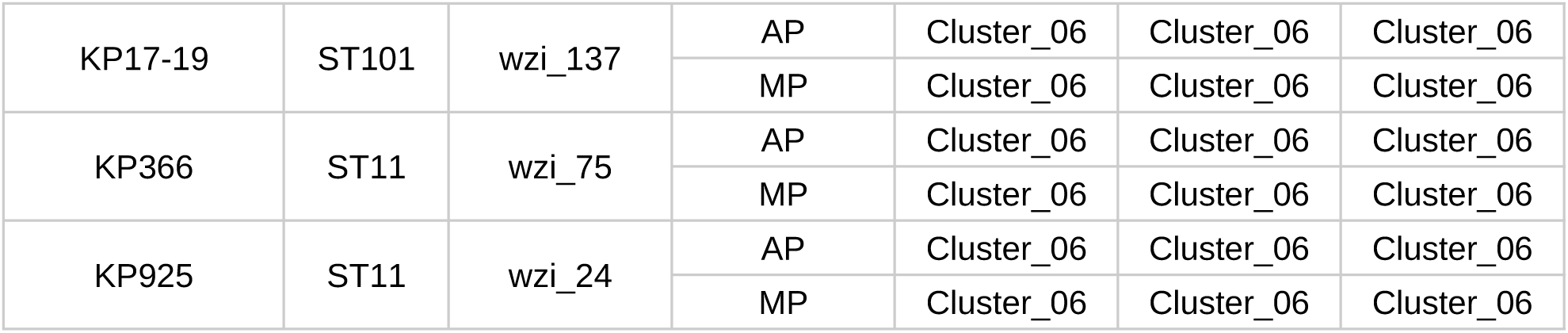
Results of clustering analysis. For each strain included in the study the cluster assigned by MeltingPlot tool on the basis of *wzi-3* and *wzi-4* melting temperatures obtained by each instrument and operator (“CFX96” for Biorad CFX96; “Eco_RT” for Illumina Eco Real-Time; “QS_Flex6” for Applied Biosystem QuantStudio 6 Flex) are reported. The relative Multi Locus Sequence Typing (MLST) profiles and *wz*i alleles (*wzi*) information retrieved from Perini and colleagues (Perini, Piazza, et al., 2020) are also reported.

| Isolate | MLST | wzi | Operator | CFX96 | Eco_RT | QS_6Flex |
| --- | --- | --- | --- | --- | --- | --- |
| 52BG | ST10 | wzi_95 | AP | Cluster_01 | Cluster_01 | Cluster_01 |
|  |  |  | MP | Cluster_01 | Cluster_01 | Cluster_01 |
| KP485 | ST15 | wzi_89 | AP | Cluster_02 | Cluster_02 | Cluster_02 |
|  |  |  | MP | Cluster_02 | Cluster_02 | Cluster_02 |
| BG-Kpn-22-18 | ST307 | wzi_173 | AP | Cluster_03 | Cluster_03 | Cluster_03 |
|  |  |  | MP | Cluster_03 | Cluster_03 | Cluster_03 |
| KP15-19 | ST307 | wzi_173 | AP | Cluster_03 | Cluster_03 | Cluster_03 |
|  |  |  | MP | Cluster_03 | Cluster_03 | Cluster_03 |
| 54BG | ST258 | wzi_29 | AP | Cluster_04 | Cluster_04 | Cluster_04 |
|  |  |  | MP | Cluster_04 | Cluster_04 | Cluster_04 |
| 56BG | ST258 | wzi_29 | AP | Cluster_04 | Cluster_04 | Cluster_04 |
|  |  |  | MP | Cluster_04 | Cluster_04 | Cluster_04 |
| BG-Kpn-21-18 | ST512 | wzi_154 | AP | Cluster_05 | Cluster_05 | Cluster_05 |
|  |  |  | MP | Cluster_05 | Cluster_05 | Cluster_05 |
| BG-Kpn-23-18 | ST512 | wzi_154 | AP | Cluster_05 | Cluster_05 | Cluster_05 |
|  |  |  | MP | Cluster_05 | Cluster_05 | Cluster_05 |
| KP359 | ST149 | wzi_62 | AP | Cluster_05 | Cluster_05 | Cluster_05 |
|  |  |  | MP | Cluster_05 | Cluster_05 | Cluster_05 |
| BG-Kpn-101-19 | ST15 | wzi_24 | AP | Cluster_06 | Cluster_06 | Cluster_06 |
|  |  |  | MP | Cluster_06 | Cluster_06 | Cluster_06 |
| KP1-19 | ST101 | wzi_137 | AP | Cluster_06 | Cluster_06 | Cluster_06 |
|  |  |  | MP | Cluster_06 | Cluster_06 | Cluster_06 |
| KP17-19 | ST101 | wzi_137 | AP | Cluster_06 | Cluster_06 | Cluster_06 |
|  |  |  | MP | Cluster_06 | Cluster_06 | Cluster_06 |
| KP366 | ST11 | wzi_75 | AP | Cluster_06 | Cluster_06 | Cluster_06 |
|  |  |  | MP | Cluster_06 | Cluster_06 | Cluster_06 |
| KP925 | ST11 | wzi_24 | AP | Cluster_06 | Cluster_06 | Cluster_06 |
|  |  |  | MP | Cluster_06 | Cluster_06 | Cluster_06 |

**Figure 2:**
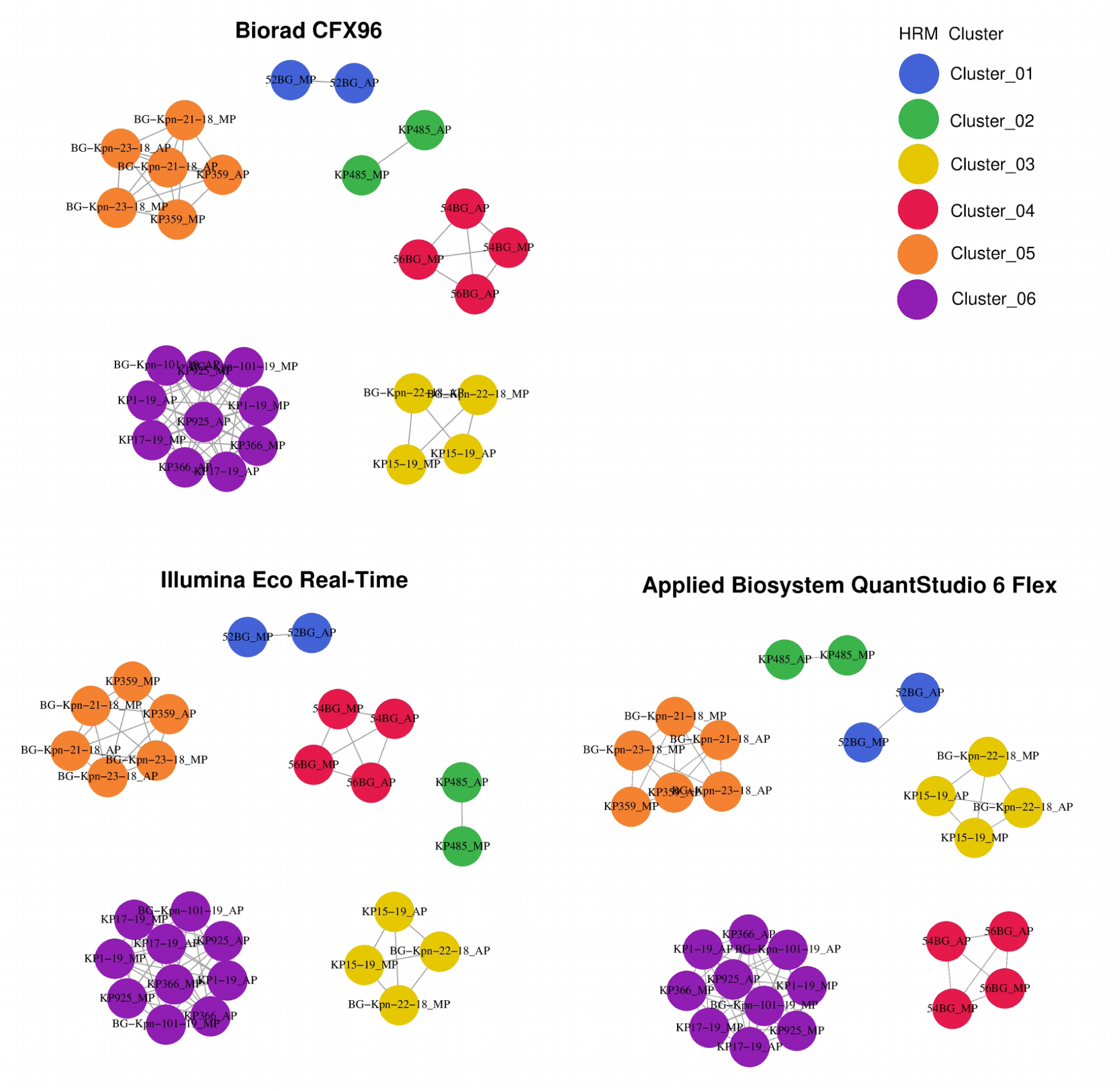
HRM-based strains clustering. The outputs of the MeltingPlot tool on the basis of *wzi-3* and *wzi-4* melting temperatures obtained by each of the three instruments included in the study (CFX96, ECO_RT and QS_6Flex) are shown. Two strains are connected if the average melting temperatures (aTm) for both wzi-3 and wzi-4 do not differ more than 0.5°C. The strains clusters were identified by MeltingPlot tool on the basis of graph topology and highlighted by different colors.

## Discussion

In the present study we evaluated the repeatability and reproducibility of the *wzi* HRM protocol for *Klebsiella pneumoniae* typing (Perini, Piazza, et al., 2020). For this validation study, we selected a subset of 14 strains representative of the entire collection of 82 *K. pneumoniae* isolates typed by Perini and colleagues (Perini, Piazza, et al., 2020). The dataset strains belong to eight different Multi Locus Sequence Typing profiles, including the most epidemiologically relevant ones (David et al., 2019; Wyres et al., 2019), i.e. ST258, ST512, ST307, ST11, ST101 and ST15. In this study we validated the protocol on three real-time PCR/HRM instruments: Biorad CFX96, Illumina Eco Real-Time and Applied Biosystem QuantStudio 6 Flex. The three instruments were placed in three different laboratories in two cities. We also studied the effect of different operators on the results. Two operators (AP and MP) independently performed HRM analysis on the 14 *K. pneumoniae* strains on the three instruments.

Statistical analyses revealed that the operators do not affect the measured melting temperatures. This results shows that the HRM protocol proposed by Perini and colleagues (Perini, Piazza, *et* al., 2020) is highly repeatable and thus reliable for large scale studies, even if several operators are involved.

Conversely, the instruments resulted to significantly affect the measured melting temperatures (Supplementary Table 5). As shown in Table S4, for some strains the difference of melting temperatures among the instruments exceeds 0.5°C, the value previously set by Perini and colleagues (Perini, Piazza, et al., 2020) as the threshold to discriminate among clusters. This result shows that strain clustering analysis can only be performed using melting temperatures obtained from the same instrument. Nevertheless, the strains clusters obtained using the MeltingPlot tool (Perini, Batisti Biffignandi, et al., 2020) were perfectly conserved among the three instruments (Figure 2 and Table 1). These results show that the melting temperature measurement process is not reproducible on different instruments. Despite this, the MeltingPlot (Perini, Batisti Biffignandi, et al., 2020) clustering analysis allows to compare the results obtained from different instruments/laboratories.

## Acknowledgements

Thanks to the Romeo ed Enrica Invernizzi Foundation.

## Supporting information

**Supplementary Table 1. Strains genomic information**

The Multi Locus Sequence Typing profiles (MLST) and *wzi* alleles (wzi) information of the strains used in this work are reported. Data was retrieved from Perini and colleagues (Perini, Piazza, et al., 2020).

**Supplementary Table 2. Instruments information**

For each instrument used in this work, the model, the short name (used in manuscript, tables and figure), the sensitivity and the location are reported.

**Supplementary Table 3. Strains melting temperatures**

The instrument used for the HRM analysis, the operator who performed the HRM analysis, the primer set used, the melting temperature replicates (T1, T2 and T3) and their average melting temperature (aTm) are reported for each strain.

**Supplementary Table 4. Melting temperature differences among instruments and operators**

The median of the average melting temperatures differences for each strain among instruments and operators are reported. In brackets, minimum and maximum difference values are reported. Maximum differences less or equal to 0.5°C are written in bold.

**Supplementary Table 5. Results of statistical analyses**

The results of statistical analyses are reported.

